# Functional Immuno-metagenomics Seizes the Alteration of Microbiota-Immune Interactions at the Early Onset of Multiple Sclerosis

**DOI:** 10.64898/2026.01.13.699307

**Authors:** Nthuku Simeon, Pons Nicolas, Parizot Christophe, Larsen Martin, Rastetter Agnes, Marie Yannick, Savine Vicart, Maillart Elisabeth, Papeix Caroline, Gorochov Guy, Imamovic Lejla

**Affiliations:** Sorbonne University, Centre for Immunology and Microbial Infections (CIMI-Paris), Inserm, Paris, France; National Institute of Agricultural Research and Environment (INRAE), University Paris-Saclay, Jouy-en-Josas, France; Department of Immunology, AP-HP Hôpital Pitié-Salpêtrière, Sorbonne University, Paris, France; Paris Brain Institute (ICM), AP-HP Hôpital Pitié-Salpêtrière, Paris, France; Department of Neurology, AP-HP Hôpital Pitié-Salpêtrière, Sorbonne University, Paris, France; Department of Neurology, Hôpital Fondation Adolphe de Rothschild, Paris Cité University, Paris, France

**Keywords:** Microbiome health, Gut microbiota, Host-Microbe Interactions, Immunoglobulin A, Immune Responses, Multiple Sclerosis, Clinically Isolated Syndrome

## Abstract

Aberrant health states are characterized by an alteration in microbial composition. Gut microbiota shapes the immune response and potential abnormal host-microbiota interactions warrant inclusion into the framework for identifying a distorted microbiota. Here, we decipher the host surveillance system based on IgA-microbe interactions and leverage it to develop the species’ Immunomodulatory score (IM_score_) to assess immunological imprint based on species’ IgA opsonisation, abundance and prevalence. We demonstrate its advantage in depicting core immunologically active gut microbiota composed of known and yet-to-be-characterized species across three independent cohorts. We then aggregated IM_scores_ into a global Immunologically active Gut microbiota index (IMG_index_) to quantify symbiotic or aberrant microbe-immune interactions at whole community level. We demonstrate IMG_index_ robustness in depicting altered microbe-immune crosstalk at the early onset of Multiple Sclerosis. Our proof-of-concept study demonstrates the importance of monitoring antibody vigilance over enteric microbes for personalised assessment of gut microbiota health.

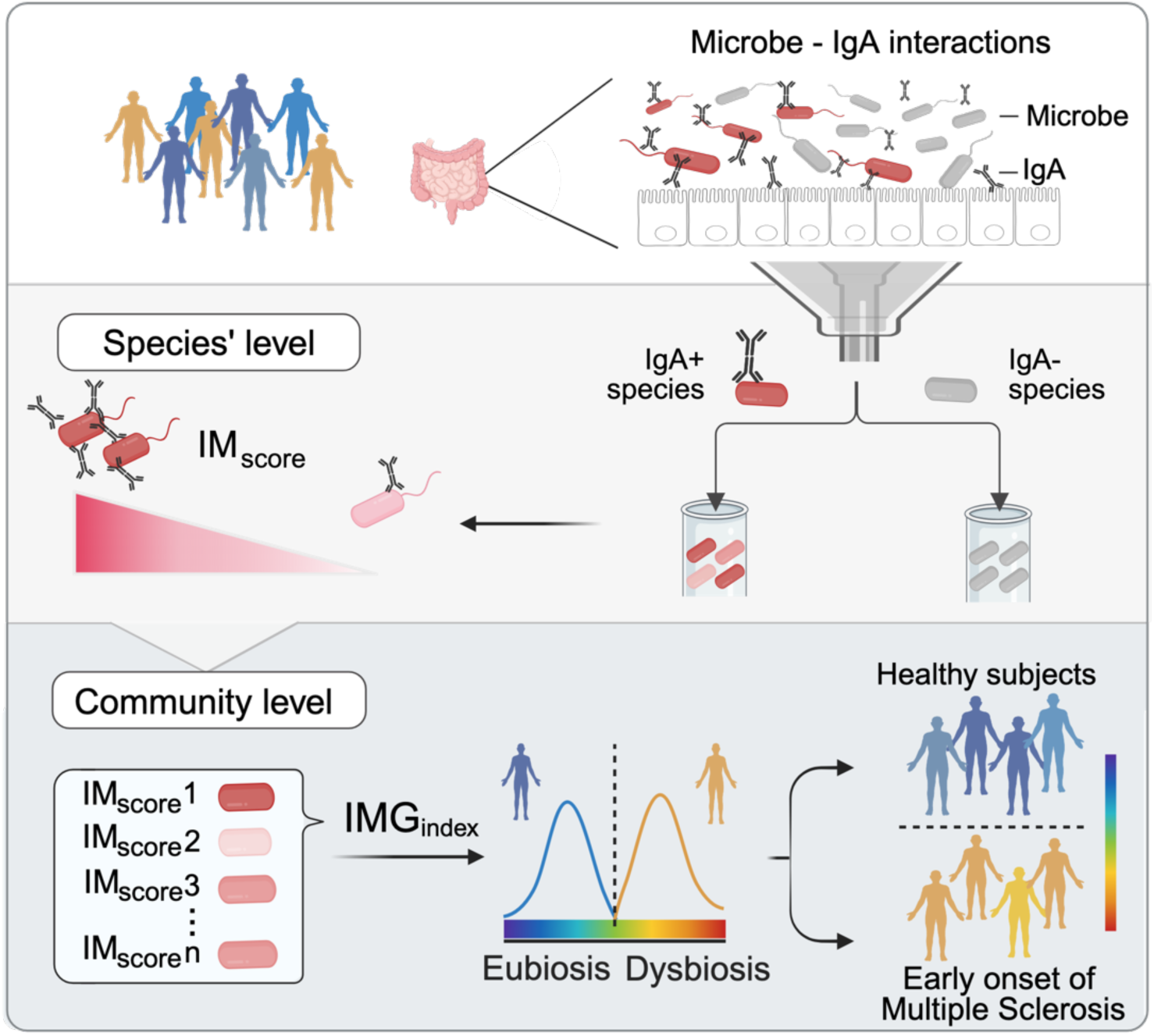

**Highlights and eTOC blurb:** Highlights

1) IM_score_ measures immunological imprint of known and uncharacterized species
2) IM_score_ highlights a core immunologically active gut microbiota
3) IM_score_ is translated into a community level IMG_index_ to measure microbiome health
4) IMG_index_ identifies aberrant microbe-immune responses at the early onset of MS

eTOC blurb

Nthuku et al. developed an approach to measure a distorted gut microbiota based on the antibody surveillance system over enteric microbes. An immunological imprint for each known and yet-to-be-characterised microbe was assessed and, then, transformed into a global community index for personal monitoring of gut dysbiosis.

## INTRODUCTION

The gut microbiome plays an important role in human health and altered microbial profiles have been observed in a wide range of diseases, including inflammatory and metabolic diseases, as well as neurodegenerative diseases such as Multiple Sclerosis (MS)^1–3^. Yet, many current approaches for identifying signs of aberrant microbiota often yield inconsistent results, as they focus on the differential analysis of single taxonomic features. To correct this bias, pioneering research has proposed the Gut Microbiota Wellness Index (GMWI) as an indicator of health status based on the abundance of selected species in the gut microbiome^4^. Recent assessments of eubiosis and dysbiosis use a concept of relational stability between microbes. This concept can be applied to identify core microbiota through taxonomic overlap, which assumes that common presence reflects biological relevance. Notably, through this lens, two recent landmark studies have identified that stably connected taxa elucidated using network analyses improve the classification of health, disease, or treatment outcomes compared to a model relying only on taxonomic composition^5,6^. Furthermore, efforts to define a health-associated gut microbiome by ranking microbial taxa according to their prevalence, stability and association with host health traits, were summarised by the Health-Associated Core Keystone (HACK) index^7^ alongside species-level ranking derived from reproducible associations with host diet and cardiometabolic health markers as proposed by the ZOE Microbiome Health Ranking^8^. Nonetheless, deployment for personalised health analysis is restricted because ecological and functional interdependencies among microbes can vary significantly depending on geographic location and individuals^9^. Moreover, such approaches do not capture the intra-kingdom interactions between microbes and the human host. Currently, despite thousands of studies, what constitutes healthy or aberrant microbiota and how host interaction with commensal bacteria affects this transition remains to be defined.

Gut microbiota is shaping the immune response by stimulating plasma cells to produce high-level secretory immunoglobulin A (IgA) at mucosal surfaces and in the gut lumen. IgA has a dual role in the gut. IgA binding to overgrowing bacteria and pathogens, prevents their growth^10^, while IgA opsonisation of commensals stimulates microbial cooperation, colonisation and production of health-associated compounds, such as Short-Chain Fatty Acids (SCFA)^11,12^. We and others previously showed that IgA recognizes diverse antigens and binds a vast number of gut microbes^13–15^. Consequently, alterations in IgA responses to gut bacteria are thought to impact susceptibility to infections and contribute to the pathogenesis of autoimmune diseases^16^. Landmark research by Berer and colleagues indicated that gut microbiota contains the factors that precipitate Multiple Sclerosis (MS), prompting research for protective and pathogenic microbial components^17,18^. MS is the most prevalent chronic inflammatory disease of the central nervous system (CNS) that potentially causes severe neurological disability^19^. In addition to genetic factors and a broad range of stimuli, gut microbiota has been implicated as a possible MS disease-modifying factor^2^. Notably, a reduction in IgA-coated faecal bacteria was observed in MS patients during relapse^16^. Similarly, we detected that the proportion of IgA-opsonized bacteria is reduced in severely affected MS-disabled patients^20^. A recent study aimed to characterise the IgA-opsonised gut microbiota in MS subjects^21^. Yet, the extremely high variations in IgA opsonisation of gut microbiota reported in the study cannot be explained by variation in microbiome composition that remain relatively modest between patients and healthy controls, and even more so between MS severity groups^2^. Therefore, despite these efforts, existing methods based on quantification remain inadequate to capture the intricate IgA-microbiota interactions.

IgA binding to gut bacteria might be impacted by species’ significant genomics diversity, including metabolic and virulence traits that are not captured using limited resolution of 16S amplicon sequencing^22,23^. Moreover, many bacteria in the gut remain unknown and the immunostimulatory contribution of this so called “microbial dark matter” remains to be elucidated. Recently, two studies simultaneously reported a method to depict higher resolution of IgA-bound bacteria in healthy subjects^22,23^. While IgA exhibits a remarkable control of gut microbiota, and *vice versa*, these dual interactions have not yet been employed as an indicator of health status, largely because immunologically active members of gut microbiota are not well characterised and, thus, are currently overlooked.

Here, we build a novel framework to decipher and quantify the host surveillance systems over gut microbiota mediated by IgA. We based our approach on the consideration that not all enteric microbes are equally targeted by IgA^13,14^. Probability of each species to be opsonised by IgA is classically calculated as an index based on species abundance in IgA+ relative to IgA-fraction^24^. However, this consideration alone is not sufficient to account for species contribution to overall immunological imprint on gut microbiota. For instance, if two species are equally targeted by IgA in terms of levels and proportions of opsonisation, the contribution of the more abundant species on overall microbiota-IgA interactions will be excessively higher. Moreover, a shared IgA-bound microbiota across the population might not only indicate a resilience to environmental impact, but also an improved ability to form stable interactions with IgA and other gut bacteria^12^. This stable relational dynamic thus relies on highly prevalent (core) microbiota members to maintain consistent functional relationship with the gut ecosystem and preserve the ecological balance^5,25^. Thus, the contribution of prevalent IgA-bound species to microbiota immunological activity is presumed to be higher. To account for this ecological perspective and measure the contribution of each taxon to overall immunological activity of the gut microbiota, we established an Immunomodulatory score (IM_score_) that merged species’ IgA opsonization levels with abundance and prevalence among the studied population (Figure 1). Using our approach, we demonstrate that numerous known and yet-to-be-characterised microbes are immunologically active members of gut microbiota. Furthermore, we show that the IM_score_ can be used to compute and quantify functional immunological impact of overall IgA-targeted gut community. Lastly, because autoimmune diseases, such as MS, are associated with changes in gut microbiota^2,3,20,26^, we demonstrate the utility of this concept in improving the distinction between healthy volunteers and treatment-naïve patients before the onset of chronic MS disease.

**Figure 1.**
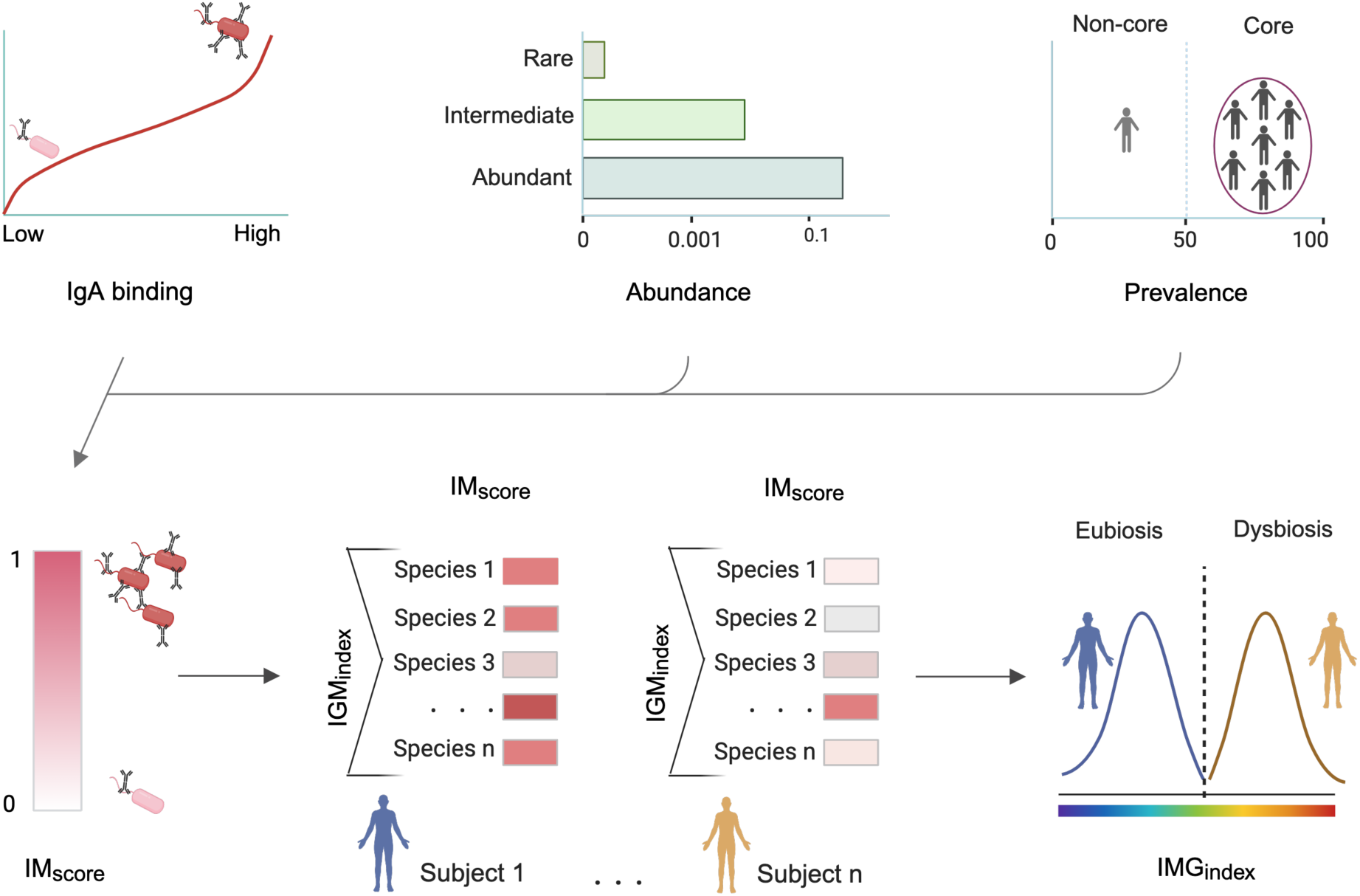
Design rationale to compute IM_score_ and <u>IM</u>munologically active <u>G</u>ut microbiota <u>index</u> (IMG_index_). To depict the species’ contribution to microbiota-IgA interactions, we develop an aggregated IM_score_ that is composed of three sets of weights: IgA binding probability, abundance, and prevalence. IM_score_ is based on high-resolution species-level taxonomic annotation of IgA-sorted microbial fractions. The IM_score_ can be further employed to compute the aggregated IMG_index_ for the overall strength of microbiota-IgA interactions for each subject. The IMG_index_ quantifies the net effect of microbiome-immune interactions on the host in gut microbiota, specifically in eubiosis and dysbiosis. As a proof-of-principle in this study, we measured the IMG_index_ in healthy subjects and patients before the onset of chronic autoimmune disease.

## RESULTS

### Known and uncharacterized microbes are the backbone of immunologically active gut microbiota

To identify the immunologically active gut microbiota, we employed shotgun metagenomic Immunoglobulin sequencing (MIG-Seq) and identified IgA-targeted microbes at species-level resolution ^22,23^. For each subject, one million IgA-bound (IgA+) and 9 million IgA-unbound (IgA-) microbes were purified using Fluorescence-Activated Cell Sorting (FACS) (Figure 2A). Sorted fractions from 18 healthy volunteers were shotgun sequenced. IgA+ and IgA-microbiota compositions were assessed using MetaPhlAn4, which allows identification of known and uncharacterized microbiota that are currently only represented by Metagenome-Assembled Genomes (MAGs) and reconstructed as Species Genome Bins (SGB)^27^. To evaluate the completeness of microbial richness represented by our IgA+ and IgA-samples, we performed rarefaction analysis of microbial diversity at different sequencing depths. We observed that rarefaction curves plateaued at 2.5M reads, indicating that most of the microbial diversity was captured (Figure S1A). To better understand the community structure and novelty of taxonomically diverse IgA-targeted microbes, we assessed whether taxa identified are known or uncharacterized species (“dark microbial matter”). These dark microbes are defined by their classification into putative taxonomical ranks as not yet fully identified species^27^. We observed that out of 455 IgA-targeted MAGs, only 55 % were known at the species level (Figure 2B), while the remaining MAGs were annotated at a higher taxonomical level and showed varying relative abundance (Figure S1B). In some subjects, these uncharacterized microbes occupied up to 53 % relative abundance of IgA-targeted gut microbiota (mean 19 %) (Figure 2C). Interestingly, 13 % of MAGs were assigned at the family level and 27 % only at the phylum level (Figure 2B) confirming the suitability of our approach to functionally characterise known and uncharacterized immunologically active microbiota.

**Figure 2.**
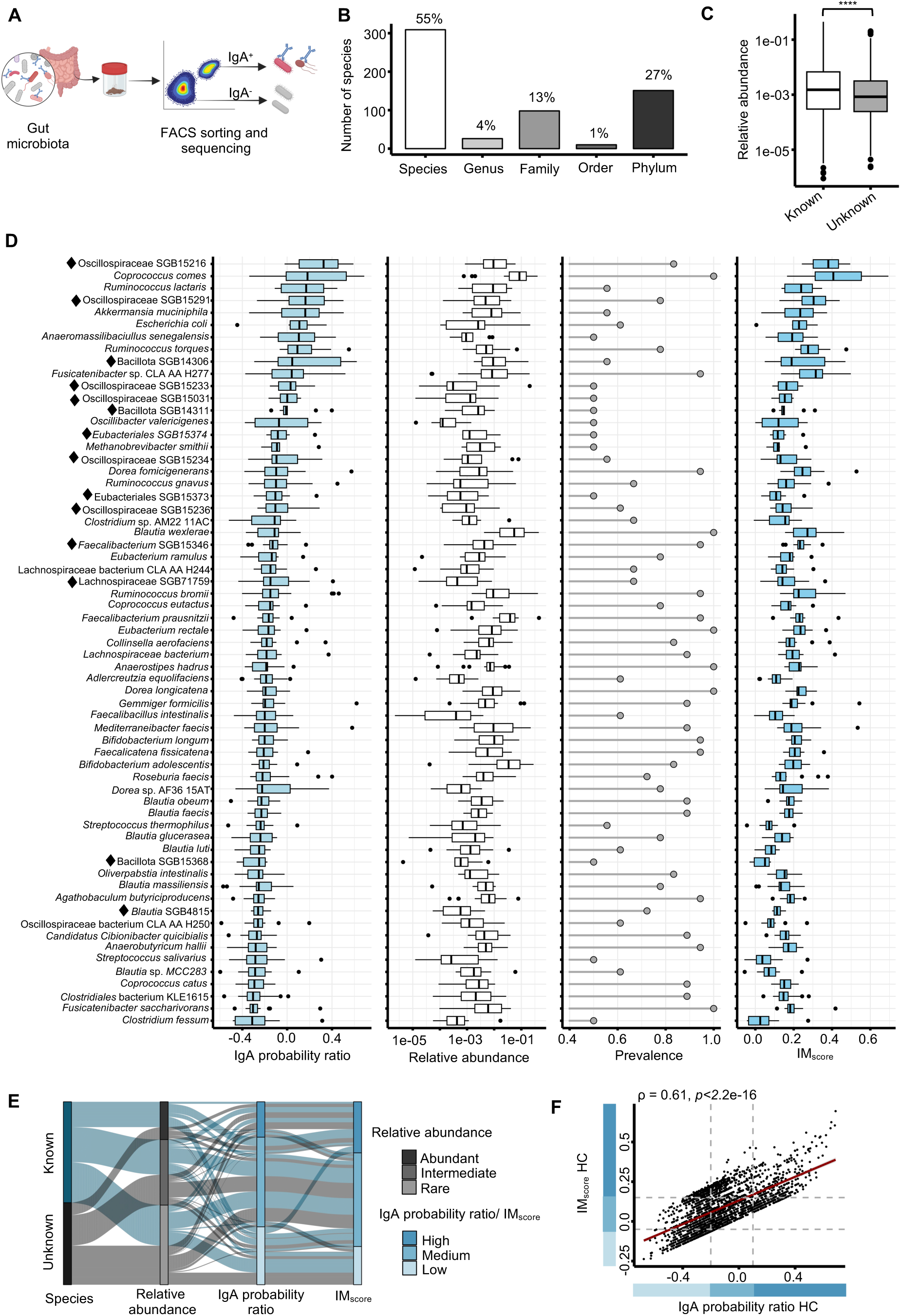
Immunomodulatory gut microbiota shows large variation in abundance and prevalence in healthy subjects. (A) High-resolution immuno-metagenomics IgA sorting workflow relies on sorting IgA+ and IgA-fractions of the gut microbiota, followed by shotgun sequencing and data analysis. (B) Number and percentage of species characterized is at indicated at each taxonomical level. Species identified only at genus or higher taxonomical level are considered as yet-to-be-characterised species. (C) Relative abundance of known and unknown (yet-to-be-characterised) IgA-opsonised species. (D) Variation of IgA probability ratio, relative abundance, prevalence and IM_score_ for core microbiota (>50% prevalence). ♦ denotes yet-to-be-characterised species. The boxplots line depicts the median value. (E) Sankey diagram representing the overall trend of known and unknown species in different categories of IgA probability ratio, relative abundance and IM_score_. (F) Spearman correlation coefficient (ρ) between IgA probability ratio and IM_score_.

We inferred the level of IgA binding of known and uncharacterized taxa using the IgA probability ratio which is calculated based on the ratio of a species’ abundance in the IgA+ vs. IgA-fractions^24^. Inclusion of IgA-unbound fraction instead of pre-sorted “native” fraction allows more precise distinction of the relative binding of microbial taxa with high and low levels of IgA binding because both IgA+ and IgA-fractions are subjected to the same workflow (*e.g*., FACS sorting conditions). According to the differences observed in IgA opsonization patterns, we group IgA-opsonised microbiota into three categories: highly targeted (IgA probability ratio > 0.1), medium (between –0.2 and 0.1) and low (<-0.2) (Figure 2D). To evaluate the community structure of IgA-targeted microbiota, we employed ecological and evolutionary constraints and classified species based on: *i*) prevalence into core (prevalence >50% across all samples) and non-core (<50%), and *ii*) abundance into abundant (relative abundance >0.1%), intermediate (0.001% – 0.1%) and rare (<0.001%) (Figure 2E). We depicted 67 core species, out of which 13 were uncharacterized and identified as members of Oscillospiraceae family SGB15216, SGB15219, SGB15233, SGB15031, *Faecalibacterium* SGB15346, or as members of Bacillota phylum (SGB14306, SGB14311) (Figure 2D). Remarkably, we observed that out of 13 species with IgA probability ratio >0, six were members of dark microbiota, confirming that these yet-to-be-characterised microbes were highly opsonised by IgA (Figure 2D, E, S1C-F). This demonstrates that microbiome contains prevalent immunologically active species that were neglected or missed in previous studies.

### Prevalent and rare microbes among immunologically active gut microbiota

To deduce the contribution of each species to immunologically activity of microbiome, we developed the IM_score_ based on three sets of features (IgA binding, prevalence and abundance of each species in IgA+ fraction) which, using a weighted sum formula, were summarized into one score (Figure 1). The IM_score_ weights were derived using a hybrid data-driven and biologically informed approach, where an Automatic Democratic Method^28^ optimized taxon contributions and regression-derived coefficients emphasized IgA binding as the dominant feature. The Automatic Democratic Method is an objective approach to generate weights that involves inferring the optimal score for each feature after which weights are deduced by regression. These data-driven weights were then biologically adjusted to prioritize IgA binding, reflecting its stronger functional relevance to host immune recognition compared to abundance or prevalence. To investigate the stability of our final weights, we performed sensitivity analysis across a range of plausible weighting combinations top-k species overlap and Kendall’s concordance. We observed high concordance among different weights (W=0.947, *p*=2.2e-16) (Figure S1G, S1H), indicating the species ranking was robust.

Next, we compared IM_score_ and IgA probability ratio moderate positive relationship between IM_score_ and IgA probability ratio was also depicted with Spearman coefficient analysis (R=0.61, *p*<2.2e-16) (Figure 2F). Given that IgA-targeted microbes had large variation in abundance and prevalence (Figure 2D, S1C-F), the IM_score_ allowed to incorporate also for species’ ecological contributions to overall IgA-interactions (Figure 2E). For instance, *Coprococcus comes*, *Ruminococcus lactaris* and *Akkermansia muciniphila* are highly opsonised by IgA. However, given that both *Ruminococcus lactaris* and *Akkermansia muciniphila are* less abundant than *Coprococcus comes* (*p*=3.74e-04 and *p*=4.84e-04, respectively), their impact on IgA interactions is reduced as reflected in lower IM_score_ value (*p*=4.85e-03, *p*=1.63e-03, respectively). These data indicate that our approach introduces an additional layer of ecological adjustments for in-depth assessment of the impact gut microbes have on the immune system.

### Dark microbiota species are also found among prevalent IgA targets in two independent cohorts

Puzzled by the high number of known and uncharacterized immunologically active microbes with a high IM_score_ (Figure 2D, E, S1E, F), we decided to validate our results using independent cohorts. Characterisation of IgA microbiota is a labour-intensive workflow and, currently, there are only two other studies that employed a shotgun approach that allows in-depth characterisation of IgA-bound microbiota. These studies included healthy individuals: 32 in the US and 10 in the Netherlands (NL)^22,23^. We retrieved the raw data from these studies, re-annotated species against the MetaPhlAn4 database^27^ and computed the IgA binding and IM_score_. Like in our own dataset, we observed a large fraction of uncharacterized, IgA-opsonised species from these two independent cohorts (Figure 3A). Remarkably, 25 out of 55 yet-to-be-characterized species that were found to overlap in the different cohorts were only characterised at the family level (20 of which are assigned to Oscillospiraceae), followed by 20 at phylum, six at genus and four at order level (Figure 3A). Notably, we observed several uncharacterized taxa among highly prevalent immunologically active microbiota, such as Oscillospiraceae SGB15216, *Faecalibacterium* SGB15346 and three species which were only annotated at the phylum level (Bacillota SGB4303, SGB15368 and SGB14969) (Figure 3A). To assess the robustness of our approach to depict these “dark” microbes targeted by IgA, we assessed the quality of reconstructed microbial genomes from IgA+ metagenomes. We observed that yet-to-be-characterised MAGs had high levels of genome completeness and low levels of contamination (Figure S2A) which further validated our approach to compute contribution of “dark” microbiota to immunological activity of gut microbiota.

**Figure 3.**
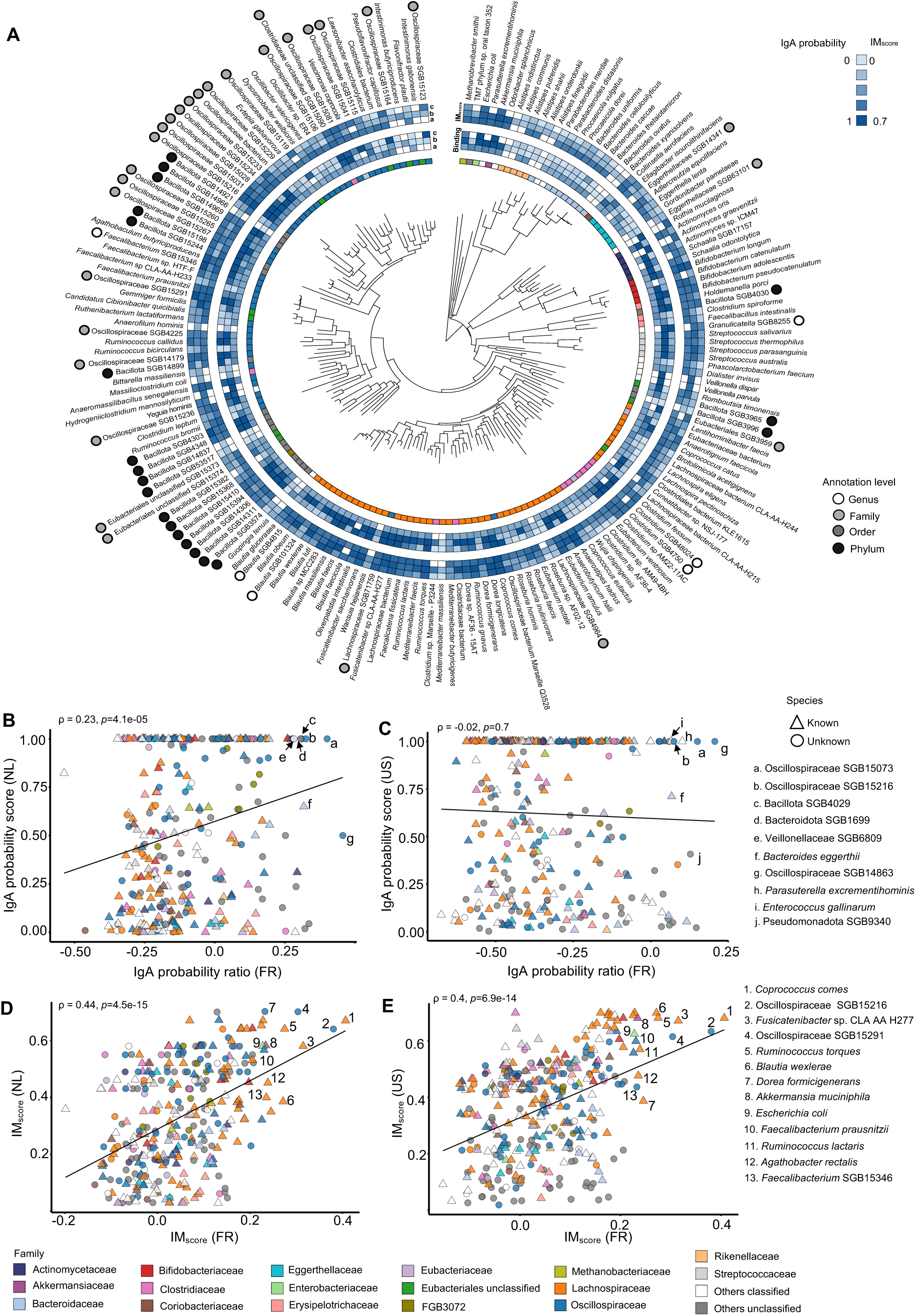
IgA binding and immunomodulatory pattern of characterized and uncharacterized species are shared across geographically distinct regions. (A) Overlap in immunologically active gut microbiota in three independent cohorts. A phylogenetic tree of species with >50% prevalence in a) France (FR; this study), b) the US (Olm *et al*.^22^), c) and NL (van Gogh *et al*.)^23^. The inner rings represent: the family of the respective species, IgA binding and IM_score_. The annotation level of yet-to-be-characterized species is indicated using the grey scale circles. (B-C) Spearman correlation (ρ) between IgA binding in FR and NL (B), and FR and US(C). (D-E) Spearman correlation between IM_score_ in our cohort FR and NL (D), and FR and US (E). Outlined top-14 species with high IM_score_.

### IM_score_ allows the definition of a core immunologically active microbiota across three cohort studies

We then computed the IM_score_ and performed an overall correlation analysis between different cohorts using IgA binding and IM_score_. Notably, when relying on IgA probability-based scoring indices, we observed a weak positive correlation (ρ=0.23, *p*=4.1e-05) between subjects in our study and van Gogh *et al.*^23^ (Figure 3B) and no correlation (ρ=-0.02, *p*=0.7) between subjects in our study and Olm *et al*. ^22^(Figure 3C). However, when comparing immunologically active microbiota in our and in other cohorts using the IM_score_, the outliers in the correlation data were reduced and we observed a moderate positive correlation between our subjects in our study in both van Gogh *et al.*^23^ (ρ=0.44, *p*=4.5e-15) (Figure 3D) and Olm *et al*.^22^ (ρ=0.4, *p*=6.9e-14) (Figure 3E). These analyses further confirm the applicability of our approach to highlight the dominant IgA response to gut microbes. Importantly, positive correlation patterns among immunologically active microbiota from different experimental set up (*e.g.,* FACS-sorting employed in our study *vs.* magnetic separation of IgA+ fractions employed by Olm *et al.*^22^ and using pre-sorted *vs.* IgA-fraction for inferring IgA binding^22,23^) and different geographic locations exemplify the robustness of functional host-microbe interactions under a variety of environmental exposures. Moreover, the overlap in IgA-bound microbiota shared across geographically distinct regions suggests that these species are part of core immunologically active microbiota that sustain IgA-microbiota symbiosis despite systematic biases due to batch effect and cofounding factors.

### Perturbed crosstalk between gut microbiota and IgA in CIS subjects unveiled by IM_score_ approach

To assess whether IgA-targeting of microbes can identify early gut dysbiosis, we sorted IgA+ and IgA-microbiota from 14 subjects from subject untreated Clinically Isolated Syndrome (CIS) (cohort characteristics are summarized in Supplementary Table 1). CIS is a condition at the early onset of MS disease^29^. We observed no difference between percentage of IgA-opsonized microbiota in HC and CIS subjects (*p*>0.05; Figure 4A). IgA-bound microbiota in HC and CIS was dominated by the members of the Lachnospiraceae, Oscillospiraceae and Bifidobacteriaceae families, followed by four uncharacterized families from the phylum Bacillota (Figure S3A). This data was consistent with previous studies that Lachnospiraceae and Oscillospiraceae are the most routinely IgA-targeted taxa in the human gut^22,23^. *Coprococcus comes*, *Blautia wexlerae*, *Faecalibacterium prausnitzii*, *Bifidobacterium adolescentis* and *Ruminococcus bromii* were the most dominant species that occupied more than 1/3 of IgA-bound microbiota (Figure 4B). We found no difference in IgA-bound microbiota α– and ß-diversity between HC and CIS (Figure 4C, D). Nonetheless, ß-diversity analysis based on Bray-Curtis dissimilarity revealed compositional heterogeneity of the IgA+ and IgA-gut microbiota of HC (R^2^=0.11, *p*=1e-04, PERMANOVA, Figure S3A). This heterogeneity was more pronounced in CIS (R^2^=0.14, *p*=1e-04, PERMANOVA, Figure S3B). We then proceeded to investigate whether immunologically-linked scoring metrics could explain a level of variation among the HC and CIS. To achieve this, we computed the Euclidean distances based on both IgA probability ratio and IM_score_ values. We did not observe distinct sample regrouping based on IgA probability ratio (R^2^=0.09, *p*=3e-03; Figure 4E). On a contrary, the Euclidean distance inferred based on IM_score_ showed clearly marked distinction of IgA-targeted microbiota in HC *vs.* CIS while also explaining a higher source of variation (R^2^=0.3, *p*=1e-03; Figure 4F). These results underline that IM_score_ can parse the complex community to reveal differences between HC and CIS clinical phenotypes that remained unnoticed using classical approaches.

**Figure 4.**
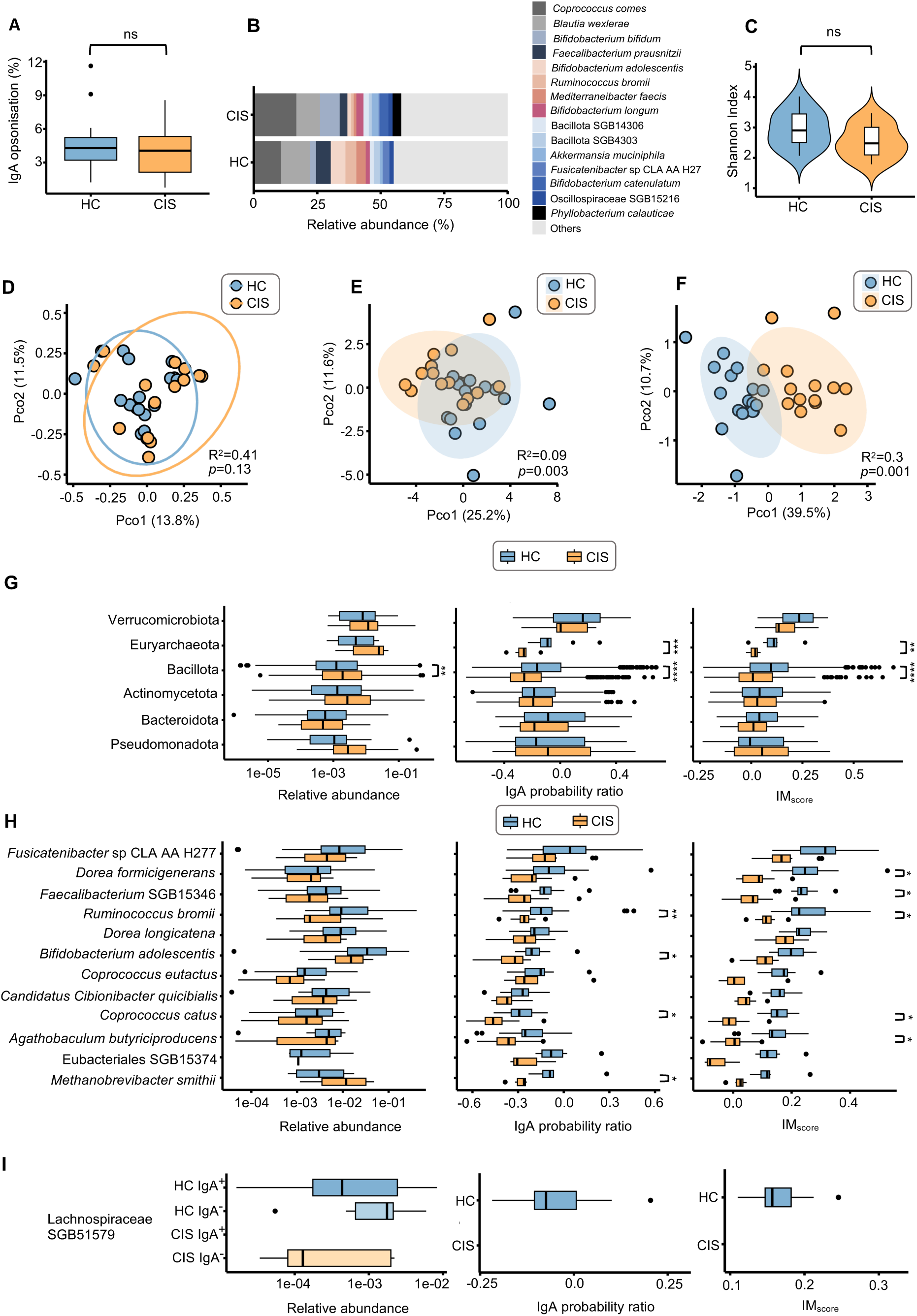
Differences in IgA-targeted microbiota between HC and CIS. (A) Percentage of IgA-opsonised microbiota in HC and CIS is measured by flow cytometry. (B) Relative abundance of the top 15 species represents half of the IgA-bound microbiota. (C) α-diversity based on Shannon index for IgA-bound species of gut microbiota in HC and CIS. (D) PCoA plot representing ß-diversity of IgA-bound species in HC and CIS groups based on Bray-Curtis’ dissimilarity. (E, F) PCoA plot based on Euclidean distances depicting the IgA probability ratio (E) and IM_score_ (F) for HC and CIS. For panels D, E and F, permutational multivariate analysis of variance (PERMANOVA) test was performed and the effect size R^2^ as well as the permutation-based *p*-value were reported. (G-I) Relative abundance (left column), IgA probability ratio (middle column) and IM_score_ (right column) at the phylum (G), species (H) and for Lachnospiraceae SGB71759 (I). (H) represents only a subset of species. Full comparison of core species depicted in Figure S3. (I) Lachnospiraceae SGB71759 was the only bacteria from HC core microbiota that was not depicted in IgA+ CIS. *p-* values for A, C, F-H were from Wilcoxon-Mann-Whitney tests with Benjamini–Hochberg (BH) correction. \**p*<0.05, \*\**p*<0.01, \*\*\**p*<0.001, \*\*\*\**p*<0.0001.

### Reduced IgA opsonisation levels of Bacillota and Euryarchaeaota in CIS

To assess the compositional differences between HC and CIS, we evaluated changes in IgA-opsonized microbiota at the phylum level. We observed increased relative abundance of Bacillota in CIS compared to HC (*p*=0.00678; Figure 4G). Contrary to this increase in relative abundance, a reduction in Bacillota interaction with IgA in CIS relative to HC was observed using both IgA probability ratio (*p*=2.03e-38) and IM_score_ (*p*=2.00e-46). We also observed a decrease in IgA-targeting of archaeal phyla Euryarchaeaota in CIS *vs.* HC (median IgA probability ratio –0.09 and –0.26 (*p*=3.33e-4) and median IM_score_ 0.11 and 0.02 (*p*=1.43e-3), for HC and CIS, respectively; Figure 4G). IgA-opsonisation of gut commensals can reduce their growth^22^, which in part could account for the lower abundance, yet higher IgA-targeting in HC *vs*. CIS depicted in this study.

### Dominant healthy immunologically active species are less abundant in CIS

We next evaluated whether there is a set of immunologically active species that are present across the HC, but altered in CIS. For this purpose, we employed ecological constraints on the species prevalence and based on this classified the taxa into core (prevalence >50 %), intermediate (prevalence 10-50 %) and infrequent taxa (prevalence <10 %). Deciphering an alteration in core microbiota in healthy subjects is relevant because one can infer the composition of homeostatic interactions of undisturbed immunologically active taxa. We observed a decrease in the number of species in CIS subjects for core microbiota in favour of an increase in relative abundance of taxa with intermediate and rare prevalence (Figure S3C), suggesting a destabilisation of IgA-targeted core microbiota that maintains immunological homeostasis.

### Microbes with reduced IM_score_ in CIS are SCFA-producing bacteria

We compared compositional alterations of IgA-bound species in CIS and observed no significant differences in overall taxa abundance (Figure S3D). We then assessed the difference in IgA-binding and depicted a decrease in IgA probability ratio for *Bifidobacterium adolescentis, Ruminococcus bromii* and *Coprococcus catus* (*p*<0.05) in CIS relative to HC (Figure 4H, S3D). In addition to bacteria, we also observed that archaeal species *Methanobrevibacter smithii* is opsonised by IgA to a lesser extent in the gut of CIS subjects relative to HC (*p*=0.0064). Notably, previous studies have depicted an increased relative abundance of *Methanobrevibacter smithii* in MS patients^30,31^. However, the impact of reduced IgA-targeting on potential expansion for this methanogenic archaeome species in MS have not been yet considered.

We then looked into the alteration of the IM_score_ in HC and CIS. Similarly, to IgA probability ratio, we observed lower IM_score_ for *Ruminococcus bromii* and *Coprococcus catus* (*p*<0.05) in CIS subjects compared to HC. In addition, a reduction in IM_score_ in CIS relative to HC was also observed for *Dorea formicigenerans*, *Agathobaculum butyriciproducens* and uncharacterized *Faecalibacterium* SGB15346 (*p*<0.05; Figure 4H, S3D). Interestingly, uncharacterized species Lachnospiraceae SGB71759 was the only species that was prevalent in HC IgA-targeted microbiota, but completely absent in the corresponding fraction of CIS patients (Figure 4I). SGB71759 was detected in the IgA-unbound microbiota in CIS (Figure 4I), indicating that the lack of IgA binding is not due to the absence of this bacterium from CIS gut microbiota. Overall, bacteria with reduced IM_score_ belong to the group of SCFA-producing microbes. Given that SCFA-producers are strong inducers of gut IgA secretion and conversely, IgA interactions with bacteria stimulate SCFA production^11^, a reduction in IM_score_ suggests a broader immunomodulatory disbalance in pre-MS subjects.

### Collapsed repertoire of species with high IM_score_ in CIS

Given the differences observed in IgA opsonization patterns for the same species within different individuals (Figure S4), we group IgA-bound microbiota into three categories based on IgA probability ratio: high, medium and low (Figure 5A). We also apply the same category for IM_score_ (high, medium and low) (Figure 5A). We then enumerated the number of species in each category and observed a larger proportion of species with high IgA probability ratio and high IM_score_ in HC relative to CIS (*p*<0.05, Chi-square test), with the difference between HC and CIS being more pronounced for IM_score_ (Figure 5A, B). Our observations are consistent with previous studies showing that IgA-targeting is highly variable between individuals ^22,23^.

**Figure 5.**
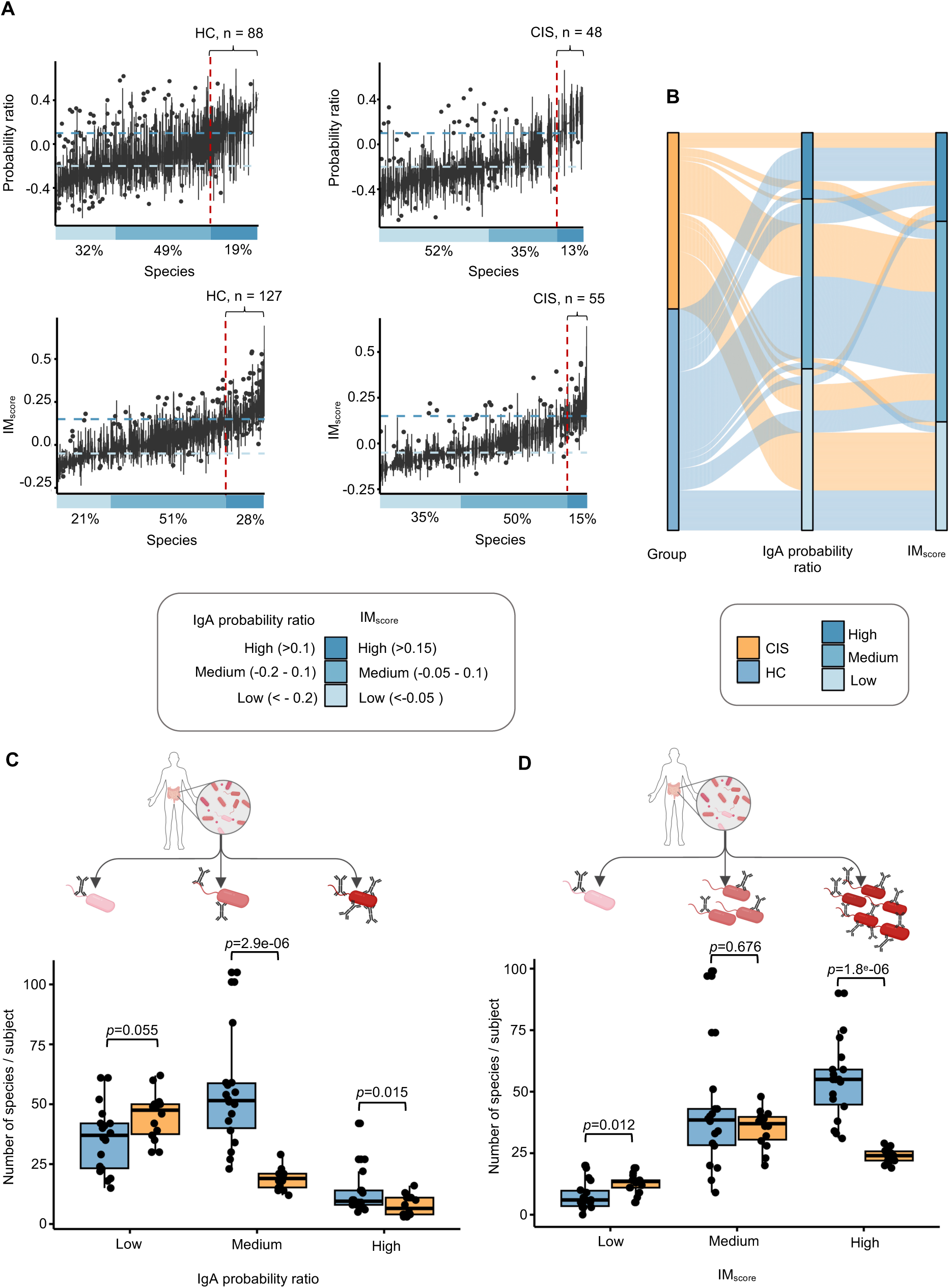
Impaired crosstalk between gut microbiota and IgA in CIS subjects. (A) Species IgA probability ratio and IM_score_ in HC and CIS. (B) Sankey diagram represents the proportions of IgA-targeted species in different categories of IgA probability ratio and IM score (high, medium, low) for CIS and HC. (C-D) IgA-targeted species in HC and CIS with high, medium and low IgA probability ratio (C) and IM_score_ (D). Category-level comparisons were statistically tested using Wilcoxon-Mann-Whitney test (BH-corrected).

To circumvent these individual variations and assess the strength of microbiota interaction with IgA in different clinical phenotypes, we enumerated the IgA-bound species in different categories of IgA probability ratio or IM_score_ (high, medium and low) for each subject. Notably, we observed a decrease in the number of taxa with high (median of 6 *vs.* 10 species per subject in CIS vs. HC, *p*=0.0147, Wilcoxon rank-sum test) and medium IgA probability ratio (median 18 *vs*. 52 species per subject in CIS vs. HC, *p*=2.9e-06; Figure 5C). The alteration was even more prominent when assessing species in high IM_score_ category. Specifically, we observed a median of 52 species with high IM_score_ in HC compared to a median of 24 species per subject in CIS (*p*=1.8e-06; Figure 5D). These results suggest a deterioration of the relationship between microbial communities and IgA in CIS compared to HC.

### IMG_index_ depicts global community perturbation of microbiota-IgA Interface in CIS

Based on the observed deterioration of IgA core microbiota in CIS relative to HC (Figure S3C), and a high individual variation in the composition of IgA-targeted microbiota (Figure S1C-F, S4), we hypothesised that only by analysing whole immunologically active microbiota as a community, we can depict stable or aberrant microbe-immune interface more optimally than when assessing the microbes individually. For this purpose, we transformed the IM_score_ that account for each species’ immunological activity based on IgA-targeting into a summarized risk index that can quantify aberrant microbe-IgA interactions at the level of the overall microbiome community (Figure 6A). We termed this <u>Im</u>munologically active <u>G</u>ut microbiota <u>index</u> (IMG_index_) and we determined it by computing the global average of the IM_score_ of all species in a sample and normalizing this average to allow for sample-based comparison (See Star Methods). We also postulated that higher IMG_index_ should indicate a homeostatic microbiota-IgA interface status. Indeed, IMG_index_ in HC maintained overall positive values, compared to mostly negative CIS values (median IMG_index_=0.468 for HC *vs.* IMG_index_=-0.716 for CIS, *p*=0.00176, Figure 6B). Importantly, once the species prevalence is established (Figure 6A), the IMG_index_ overcomes personal variation in species composition and provides an opportunity to characterize gut eubiosis or dysbiosis at the individual level.

**Figure 6.**
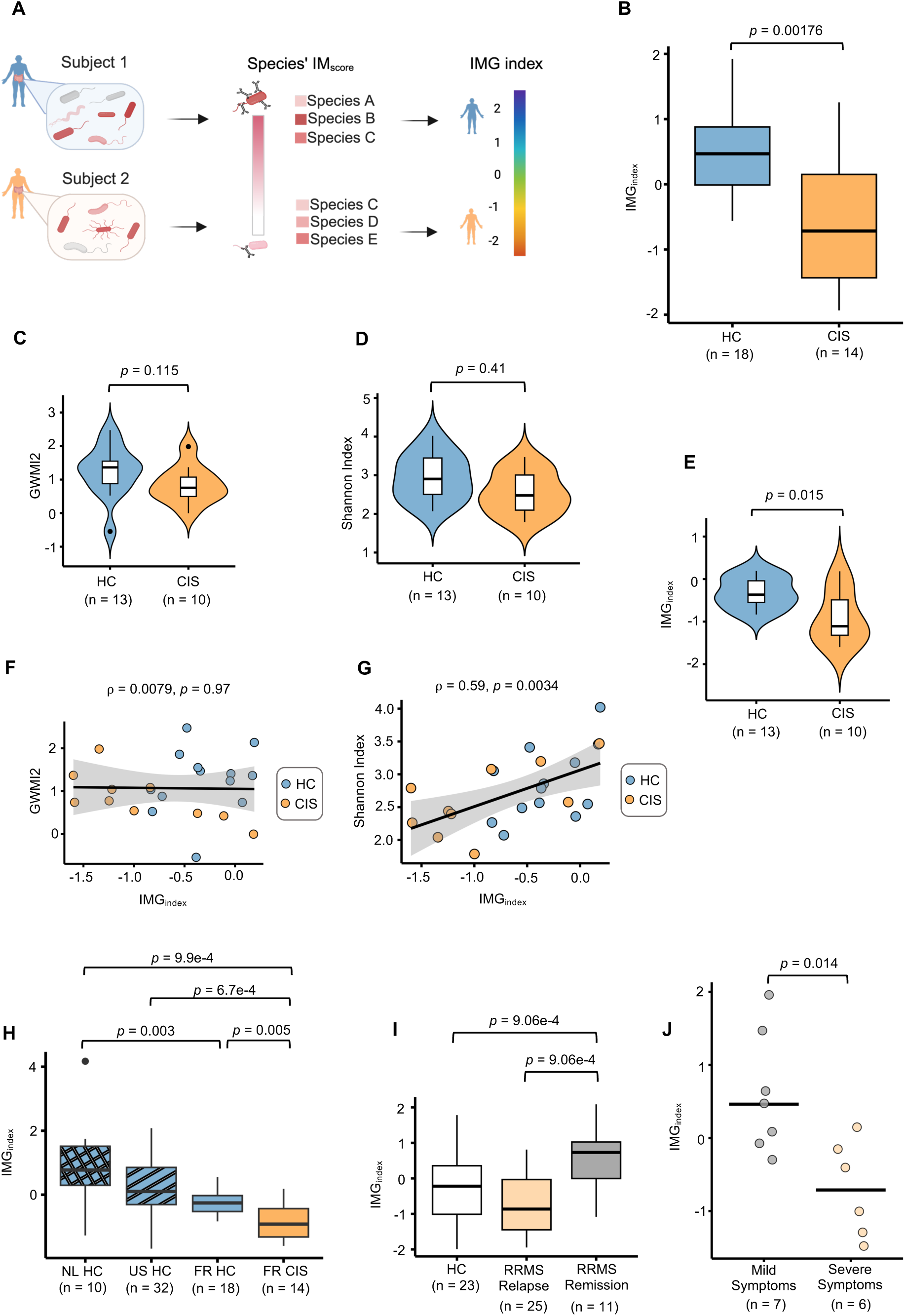
IMG_index_ captures deterioration of gut microbiota homeostatic interaction with IgA. (A) Schematic illustration for rationale beyond IMG_index_ calculation for HC and CIS subjects. (B) Differences in IMG_index_ for HC and CIS subjects. (C-E) GMWI2 (C), Shannon index (D) and IMG_index_ (E) for HC and CIS subjects. (F-G) Spearman correlation coefficient between (π) IMG_index_ and GMWI2 (F), between and IMG_index_ and alpha diversity of whole microbiota (Shannon index) (G). (H-J) Validation of IMG_index_ using publicly available dataset for IgA-bound microbiota. (H) Higher IMG_index_ for heathy individuals (HC) across three different regions (NL^23^, the US^22^, and France (FR, this study) was depicted in all HC groups relative to CIS subjects in this study. (I) Variation of IMG_index_ in RRMS patients corresponds to remission and clinical relapses during active MS. (J) IMG_index_ decreased in mice model with severe MS-like symptoms in comparison to mice with mild symptoms. *p*-values were obtained using Wilcoxon-Mann-Whitney with BH correction.

We next assessed the performance of the IMG_index_ compared to other similar gut microbiome health scoring approaches to depict healthy and aberrant gut microbiota. For this purpose, we obtained metagenomic shotgun sequencing data for total gut microbiota from 13 HC and 10 CIS individuals. We then computed Gut Microbiome Wellness Index 2 (GMWI2), which is proposed to predict health status based on gut microbiome taxonomic profiles^4^. We observed no difference between GMWI2 scores of HC and CIS groups (*p*=0.115; Figure 6C) nor could we observe differences in Shannon index (*p*=0.41; Figure 6D). On contrary, we revealed through IMG_index_ differences between HC and CIS patients (*p*=0.015; Figure 6E). While no correlation was observed between GMWI2 and IMG_index_ (π=0.0079, *p*=0.97; Figure 6F), a moderate positive correlation was observed between IMG_index_ and Shannon diversity (π=0.59, *p*=0.034; Figure 6G). Together these data indicate that monitoring of gut microbiota by taking into account the dimension of IgA recognition represents a powerful approach to depict dysbiosis.

### Confirmation of MS-associated decreased IMG_index_ in independent cohort and in murine model

We then proceeded to evaluate whether our results can be replicated in two additional independent healthy individual cohorts from the Netherlands^23^ and the US^22^. We observed that the IMG_index_ was higher in all healthy groups relative to the CIS (Figure 6H). To further verify the ability of the IMG_index_ to discriminate MS subjects with different disease courses, we next leveraged publicly available data on IgA-opsonised microbiota in Relapsing-Remitting Multiple Sclerosis (RRMS)^32^. RRMS is the most common form of MS that is characterised by periods of relapse and remission during which their disability is gradually getting worse^29^. While the initial study found no significant differences between IgA-bound microbial taxa in different disease states^32^, we observed an increased IMG_index_ in MS patients while in remission compared to patients during active MS corresponding clinical relapses (*p*=0.00906; Figure 6I). Additionally, we profiled IgA-targeted microbiota in MS pre-clinical mice models that developed mild and severe Experimental Animal Encephalomyelitis (EAE) ^33^. We calculated the IMG_index_ for mice with mild and severe EAE and, as expected, observed that mice with severe EAE had a lower IMG_index_ compared to mice with mild EAE (*p*=0.014; Figure 6J). This emphasizes that the IMG_index_ provide sensitive microbe-IgA co-ecology measures that confine eubiosis and dysbiosis patterns of microbiota and immune system interplay.

## Discussion

A healthy gut microbiome can be envisioned as one that maintains immunological homeostasis and stability^1^. Here we have developed the Immunomodulatory Gut Microbiota Index, a novel approach based on IgA-surveillance of microbes that can detect the aberrant gut microbiota signatures that are not observable in classical microbiome data. The recent study on IgA-microbe interaction by Olm et al., underlined IgA-bound microbes to plausibly contribute to the immune function of the gut microbiota^22^. We go beyond previous studies, including our research^13,20–23,34^, that only assessed IgA-bound microbiota through its sheer ratio to IgA-unbound microbes. With the goal to adopt an eco-evolutionary perspective, we integrated ecological and functional interdependencies between microbiota and the immune system. We developed the IM_score_ metric by considering a taxon’s relative abundance and prevalence, which permitted us to assess a microbe’s immunomodulatory properties. Moreover, by using an expanded resolution of taxonomic profiling of known and uncharacterized microbial species, we resolved a common pitfall of microbial annotations due to unstandardised annotations with current official nomenclatures. For instance, a core gut commensal and highly IgA-opsonised and abundant gut commensal, *Coprococcus comes*, in current GTDB database is represented using different names (*Coprococcus comes, Bariatricus comes*)^35^, which in spite of high-resolution shotgun sequencing, can introduce incoherent taxonomical annotations. Using the IM_score,_ we show that a previously uncharacterized, but important fraction of the gut microbiota gut bacteria, is immunologically active. In addition, we identify for the first-time immunologically active prevalent microbial taxa commonly shared among populations from different geographic regions.

Given that IgA-opsonised microbiota captures not only the structure of microbiota, but also reflects alterations in host IgA repertoire^36^, our approach reveals host/microbiota homeostasis over mere microbial taxonomy that can be employed to assess overall gut health. Applying this approach to overall community allows the assessment of the IgA-targeted microbiome as a whole community.

We show that IgA-bound gut microbes are predominantly from Lachnospiraceae and Oscillospiraceae families, which is in concordance with our and other studies on IgA-bound microbiota^13,22,23^. Many species from these families contribute important functions, such as production of anti-inflammatory SCFA that are considered a mediator in microbiota-gut-brain crosstalk^37^. In a positive feedback-loop manner, SCFA-producing bacteria (in particular, acetate producers such as *Coprococcus* sp.) are strong inducers of IgA secretion in the gut and, conversely, IgA binding to gut bacteria stimulates the SCFA production^11,38^. Our results here indicated that the IgA targeting of SCFA-producing bacteria is declining even before the disease converts into a chronic condition. This is of relevance because microbiota-produced SCFA have been shown to alleviate neuroinflammation via SCFA production^39^, and a reduction in SCFA was reported in CIS and MS^40,41^. Previous study focusing on depicting core microbial signatures underlying the enrichment of pathways related to SCFA production have found that these signatures are enriched in healthy microbiota^5^. Finally, propionic acid supplementation in MS patients was shown to reverse Treg/Th17 cell imbalance via increased Treg cell induction and enhancement of Treg cell function and was even proposed to be associated with disease course improvement^42^.

Microbial eubiosis (and dysbiosis) has gained important awareness as feature of both physical and mental health. In particular, an increasing interest in the role of microbiota in the development of autoimmune diseases raises the importance of identifying the aberrant microbial patterns even before development of chronic disease^1,2^. Yet, altered patterns in microbiome data are usually based on the data from large patient cohort and are complicated to employ at the individual level or in studies with small samples size ^4,5^. Current studies that are focused on computing healthy gut indices are focused on core microbiome is largely shared among individuals^2,4^, however, not all shared microbes are contributing equally to the immunomodulatory gut microbiota functions. In addition, focusing only on shared microbes, can lead to disregarding others, possibly critical, immunomodulatory microbes within the individual host. Yet, this wide inter-individual taxonomic diversity in composition gut microbes is often an obstacle in defining microbiome health^2,4^ and different individuals do not respond to the same microbe in the same way^43^. We encompass this variability of immunoselection by focusing on summary index the reflects the strength of microbiota interaction with antibodies. We leveraged these microbiome–immune interactions to characterize dysbiosis, supported by our observation that healthy individuals from geographically distinct regions exhibit consistent IgA opsonization patterns whereas these binding profiles are noticeably shifted in individuals with diseases, as exemplified here in the context of MS. In broader context, our approach can provide a framework to re-define the global gut microbiota health index by providing the universal strategy to study overall community over the individual species or taxonomical core microbiome structure.

## Limitations

This study highlights the potential of assessing microbial immunomodulatory activity to infer microbiome-associated health. However, our analyses were based on a subset of the microbial community, which may not fully capture the total diversity of IgA-opsonised microbiota in the human gut. Thus, our approach primarily reflects dominant IgA responses to taxa, which limits our ability to characterise IgA binding to rare or non-keystone species. Additionally, IgA targeting of pathogenic or pathobionts is not considered. Similar to other studies, despite a rigorous methodological approach being applied, the possibility of non-specific IgA binding that may introduce noise in IgA-binding profiles cannot be excluded. Lastly and importantly, the findings require validation in larger cohorts and longitudinally monitored populations to confirm their robustness and generalizability.

## Methods

### Patient enrolment and eligibility criteria

Patients with CIS and MS along with age– and sex-matched heathy individuals were enrolled. CIS patients were recruited at the Fondation Opthalmologique Adolphe de Rotschild (Paris, France) or at the Department of Neurology of Pitié-Salpêtrière Hospital (Paris, France). CIS was defined as a first central nervous system inflammatory event that lasts at least 24 hours^44,45^. The participants were adults between 18 and 61 years old without any other known immune, gastrointestinal and infectious diseases, nor previous extended intestinal surgery. The study inclusion criteria specified no use of antibiotics, corticosteroids or laxative drugs in the last three months prior the study. The study had been approved by the local ethics committee of Pitié-Salpêtrière Hospital (CPP Ile-de-France VI). A prior written consent was obtained from all the patients and controls before inclusion in the study. Detailed description of patient cohort is listed in Table S1.

Both blood and stool samples from CIS patients were collected before therapy with steroids. The study focused on isolated first clinical events, including CIS and Magnetic Resonance Imaging (MRI)-defined early MS according to the 2010 McDonald diagnostic criteria^46^. No statistical methods were employed to predetermine sample size. The experiments were not randomized and investigators were not blinded to outcome assessment.

### Sample collection and purification

Stool samples were collected in a container including a reagent for the generation of an O_2_-depleted and CO_2_-enriched atmosphere (Anaerocult band, Mikrobiologie), aliquoted in an anaerobic atmosphere and stored at –80°C. Faecal bacteria were purified by gradient purification as previously described^14^. Bacterial extracts were suspended in 1×PBS-10% glycerol, immediately frozen in liquid nitrogen and then stored at –80°C.

### Flow cytometry and isolation of IgA-coated bacterial fractions

Secretory IgA binding to microbiota was assessed by bacterial flow cytometry as previously described ^13,14^. Briefly, thawed microbial cell collected (10^8^ bacteria/condition) were suspended in 1x PBS, 2% BSA (Sigma), 0.02% Sodium azide (Sigma) and incubated in with secondary conjugated antibodies (goat anti-human IgA-PE) or isotype controls (both from Jackson Immunoresearch Laboratories) 20 minutes at 4°C in the dark. Then, bacteria were suspended in sterile PBS. One million positive bacterial fraction (IgA+) and nine million negative bacterial (IgA-) fractions were acquired as events using Fluorescence-Activated Cell Sorting (FACS) on S3 Cell Sorter (Bio-Rad). Frequencies of IgA-bound microbiota was recorded as percentages.

### DNA Extraction

Metagenomics DNA was extracted from whole stool samples as previously described^47^. Briefly, 200 mg of faecal sample was lysed chemically (guanidine thiocyanate and N-lauroyl sarcosine) and mechanically (glass beads) followed by elimination of cell debris by centrifugation and precipitation of genomic DNA. Qiagen DNeasy PowerClean Cleanup Kit was used to obtain highly pure DNA. DNA concentration was measured by One Step DNA fluorometric method on Quantas instrument (Promega). DNA from IgA+ and IgA-fraction was performed by using bead beating at max vortex speed for 12 min using Disruptor plates (Omega Biotech), followed by proteinase K treatment for 1 h at 56 C. DNA was then extracted using reagents provided with Viral DNA/RNA extraction kit (Omega Biotech).

### NGS library construction and sequencing

Shotgun NGS libraries were constructed using FX QIASEQ kit (Qiagen). Specifically, 7 µL extracted DNA was sheared enzymatically, ligated to adaptor oligonucleotides with appropriate barcodes. IgA+ and IgA-fractions were PCR amplified (12 cycles) and sequenced on NovaSeq6000 (PE500) or NextSeq2000. NGS libraries from total metagenomic DNA were sequenced directly after adaptor and barcode addition (without PCR amplification step) on NovaSeq X series.

### Sequencing data pre-processing and taxonomic classification

Raw metagenomic sequencing reads were first processed to trim adapter sequences and low-quality bases using fastp (v0.23.4)^48^. Trimmed reads were then decontaminated by removing human read pairs in which one or both aligned to the *Homo sapiens* reference genome GRCh38 using Bowtie2 (v2.5.4)^49^ Unmapped reads were then retrieved using kneaddata (v0.7.7)^50^ for downstream processes. At each step, read quality was assessed using FastQC (v.0.11.9)^51^ and summarized using MultiQC (v1.23)^52^. Quality-controlled reads were taxonomically profiled at species level using MetaPhlAn4 (v4.1.1)^27^ which harnesses a marker gene database for 26,970 species-level genome bins (SGBs). Taxonomic quantification was also performed using MetaPhlAn4 which provided relative abundance information for the metagenomic samples. Both microbial profiling and quantification were done using MetaPhlAn 4 default parameters.

### Rarefaction analysis

*In silico* rarefaction analysis was performed by subsampling six deeply sequenced IgA+ and IgA-samples (all between 20 million and 30 million reads) using bbmap reformat.sh tool ^53^. Subsampling was done at different sequencing depths after which taxonomic profiling was performed using MetaPhlan4 and species richness inferred using the Shannon diversity index.

### Inferring the binding of IgA to a microbe

The probability that a microbial taxon is bound by IgA was estimated using the IgA probability ratio and the IgA probability score^24^. The IgA probability ratio was used in the case where IgA+ and IgA-bacteria cell fraction gates sizes were available. In the absence of sorted bacterial fraction gates sizes, we employed the IgA probability score which considers the native (pre-sort) size of the bacterial cell fraction. Pseudo-count imputation was done for avoiding zero-division negation.

### Establishing the Immunomodulatory score (IM_score_)

IM_score_ was determined by considering the weighted sum of the prevalence, relative abundance and IgA binding value of a microbial taxon in a sample. The weights of the IM score were derived using a hybrid data-driven and biologically informed approach. First, we applied the Automatic Democratic Method (ADM)^28^ which is an objective approach for deriving composite scores from multiple criteria without predefined weights. For each criterion, an optimal score is calculated after which regression is performed on the optimal scores to produce coefficients that can be normalized to interpretable weights. The approach is termed “democratic” since the weights are derived collectively from the entire dataset without subjective weight specification. Here, we implemented it as follows;

For each parameter *i* that we considered, we computed an optimal score 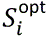 by solving a linear programming problem that maximizes a weighted sum of the three entities while enforcing a non-negativity constraint:

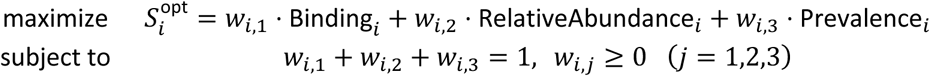

We then fit a linear regression across these parameters where the regression coefficients β_1_, β_2_, β_3_ capture the relative importance of each parameter:

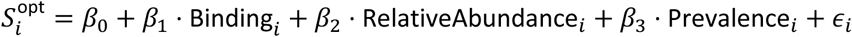

The coefficients were then normalized to one to give the final weights:

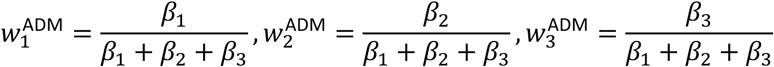

This was applied to the IgA-bound healthy control cohorts to estimate initial weights for binding probability, relative abundance and prevalence. The resulting ADM-derived coefficients indicated an inclination towards higher weights relative to IgA binding hence reflecting its dominant contribution in distinguishing microbial profiles within the healthy IgA**-**sorted samples. To ensure biological interpretability and to reflect the mechanistic relevance of microbial-host interactions, these data-driven weights were adjusted based on biological reasoning. Since microbial binding to host immunoglobulins is a more direct functional indicator of immune recognition that relative abundance or global microbial community prevalence, greater weight was assigned to the binding feature, as was backed up by the ADM method. Relative abundance and prevalence were assigned smaller and equal weight contributions to account for ecological representation and population level stability. This hybrid ADM-biological weighting approach preserved the objectivity of data-driven weights while integrating domain knowledge to ensure that the IM_score_ remains biologically meaningful and interpretable.

Hence for a taxon *i* identified in sample *j*, the IM_score_ is inferred as:

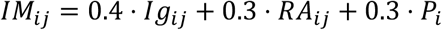

where:

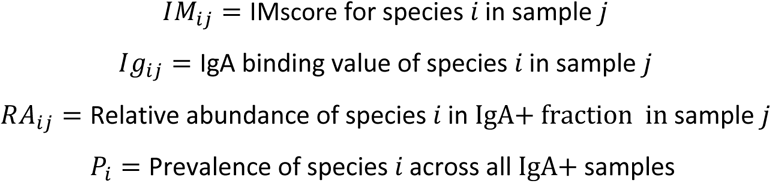

### Determining the Immunologically active Gut microbiota index (IMG_index_)

IM_score_ was used as a basis to estimate an overall community IMG_index_. To do this, the IM_score_ underwent min-max scaling to provide comparable per-species per-sample IM_score_ values.

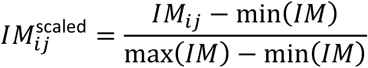

Then the global average of the scaled IM_score_ values for all species in a sample was calculated which is:

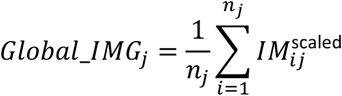

Finally, to obtain IMG_index_, the global average was standardised by performing Z-score normalization. This normalization permits comparison of how the IMG_index_ of a given sample’s deviates from the average across all samples.

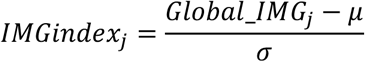

where:

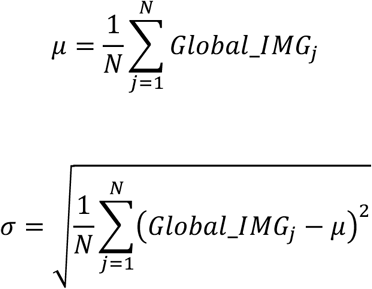

### Metagenomic assembly, binning and sequence similarity analysis

The quality-controlled metagenomic paired-end reads were assembled individually using MEGAHIT (v1.2.9)^54^ with default parameters (minimum and maximum kmer sizes of 21 and 99 respectively). Assembled contigs with a length of less than 500bp were filtered out to reduce on downstream assembly artefacts. Sample-based contigs were then binned using MetaBAT2 (v2.12.1)^55^ into microbial metagenome assembled-genomes (MAGs) where contig coverage depth was determined by mapping reads back to the filtered assemblies using Bowtie2 (v2.5.4)^49^ after which contig read depth was inferred using MetaBAT2. The quality of the resulting MAGs was assessed using CheckM2 (v1.2.3)^56^ by evaluating their genome completeness and contamination. Medium and high-quality MAGs underwent dereplication using dRep^57^ (v3.5.0) where an average nucleotide identity (ANI) of 98% was used to achieve a subspecies-level dereplication (command: comp 50 –con 10 –-S algorithm skani –-S ani 0.98 – –cov thresh 0.25 –-multiround primary clustering). Dereplicated MAGs were then classified taxonomically using GTDB-Tk (v2.4.0)^58^ to form a study-specific microbial genome database. To extract yet-to-be-characterized microbial MAGs from the dereplicated MAGs, we employed skani^59^ to measure the sequence similarity between the MAGs and yet-to-be-characterized representative genomes obtained from Pasolli et al.^60^. A dereplicated MAG was concluded to be the same species as an uncharacterized representative genome if it satisfied an ANI of >95% and an alignment fraction of > 60%.

## Statistical analysis

All statistical analyses and data visualization were performed on R (v4.2.3). Microbial alpha diversity was estimated using the Shannon diversity index while beta diversity was calculated using Bray-Curtis dissimilarity and represented using Principal Coordinate Analysis (PcoA) where statistical variance explanation was obtained using the permutational multivariate analysis of variance (PERMANOVA). Spearman’s rank correlation was used to assess the relationships between IgA probability ratio, IgA probability scores and IM_score_ values. Unpaired within-group differences were assessed using the Wilcoxon Mann-Whitney test, while paired group differences were evaluated using the Wilcoxon signed-rank test.

For analyses involving multiple comparisons, *p*-values were adjusted using the Benjamini–Hochberg (BH) false discovery rate (FDR) correction. Statistical significance was defined as *p*<0.05.

## Acknowledgments

We thank all volunteers for their participation in this study. We also thank all members of our labs for discussions as well as the Paris Brain Institute (ICM) and Institut Curie for sequencing. We thank Jun Li for his critical reading of this manuscript. This work was supported is supported by the French National Research Agency (ANR) grant JCJC ILLUVIR ANR-23-CE15-0012-01, the Foundation for Medical Research (FRM EQU202203014622), the European Innovation Council (EIC) Pathfinder NUTRIMMUNE (101162457) and the French Ministry of Health in the context of MESSIDORE 2024 call operated by IReSP (AAP-2024-MSDR-371971).

## Author contributions

Conceptualisation: S.N., G.G., L.I. Data curation: L.M., S.V., M.E., P.E., C.P., N.P., A.R., Y.M., S.N., L.I. Formal analysis A.R, M.L., S.N., L.I., N.P. Funding acquisition: G.G., L.I. Investigation: S.N., L.I. Methodology: S.N., L.I. Project administration: L.I. Resources: S.V., L.M., M.E., P.E., S.N., L.I., G.G. Software: S.N. Supervision: G.G., L.I. Validation: G.G., L.I. Visualisation: S.N., L.I. Writing – original draft: S.N., G.G., L.I. Writing review & editing: all authors.

## Tables

**Table S1.** – A description of the clinical and demographic features of the healthy donors and CIS patients.

## Supplementary Figures

**Figure S1.**
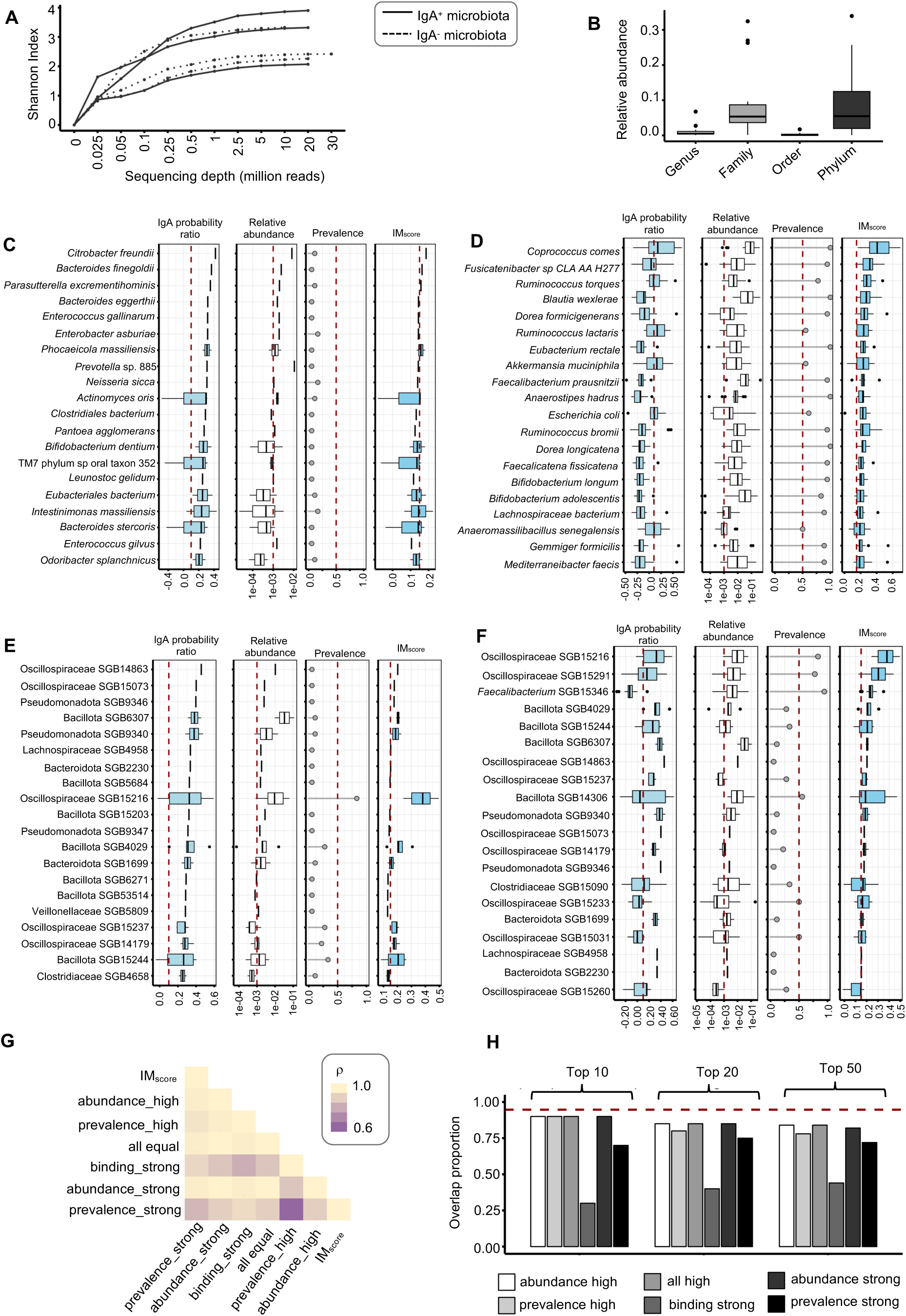
Composition of IgA-bound microbiota in healthy subjects, related to Figure 2. (A) Rarefaction curves representing species’ Shannon index against their sequencing depth. One million of IgA-bound and 9 million of IgA-unbound microbial cells were sorted with FACS and sequenced. Sequencing reads were subsampled and the Shannon index was depicted at different sequencing depth for IgA+ and IgA-microbes. (B) Relative abundance of uncharacterized MAGs annotated at the lowest taxonomical level indicated on x-axis. (C-F) Top 20 characterized and uncharacterized microbes ordered based on IgA probability ratio and IM_score_ where (C) represent the characterized microbes with the highest IgA probability ratio, (D) represents characterized microbes with the highest IM_score_, (E) shows the uncharacterized microbes with the highest IgA probability ratio and (F) shows the uncharacterized microbes with the highest IM_score_ values. (G) Spearman correlation coefficient (ρ) between the selected IM_score_ weight scheme (IgA binding, relative abundance, prevalence is weighed 0.4, 0.3, 0.3, respectively, in an aggregated score) and scores derived from additional weighting schemes. Based on the feature combination of IgA binding, relative abundance and prevalence, each additional weighting scheme emphasizes on a component feature as follows: abundance_high (0.3, 0.4, 0.3), prevalence_high (0.3, 0.3, 0.4), all equal (0.3, 0.3, 0.3), binding_strong (0.6, 0.2, 02), abundance_strong (0.2, 0.6, 0.2), prevalence_strong (0.2, 0.2, 0.6). Weighs are indicated in following order: IgA binding, relative abundance, prevalence. (H) Overlap of the top 10, 20 and 30 species across IM_score_ different weight schemes presented in (F). Kendall’s concordance coefficient depicted a high degree of agreement among weight schemes (W=0.947, red dotted line).

**Figure S2.**
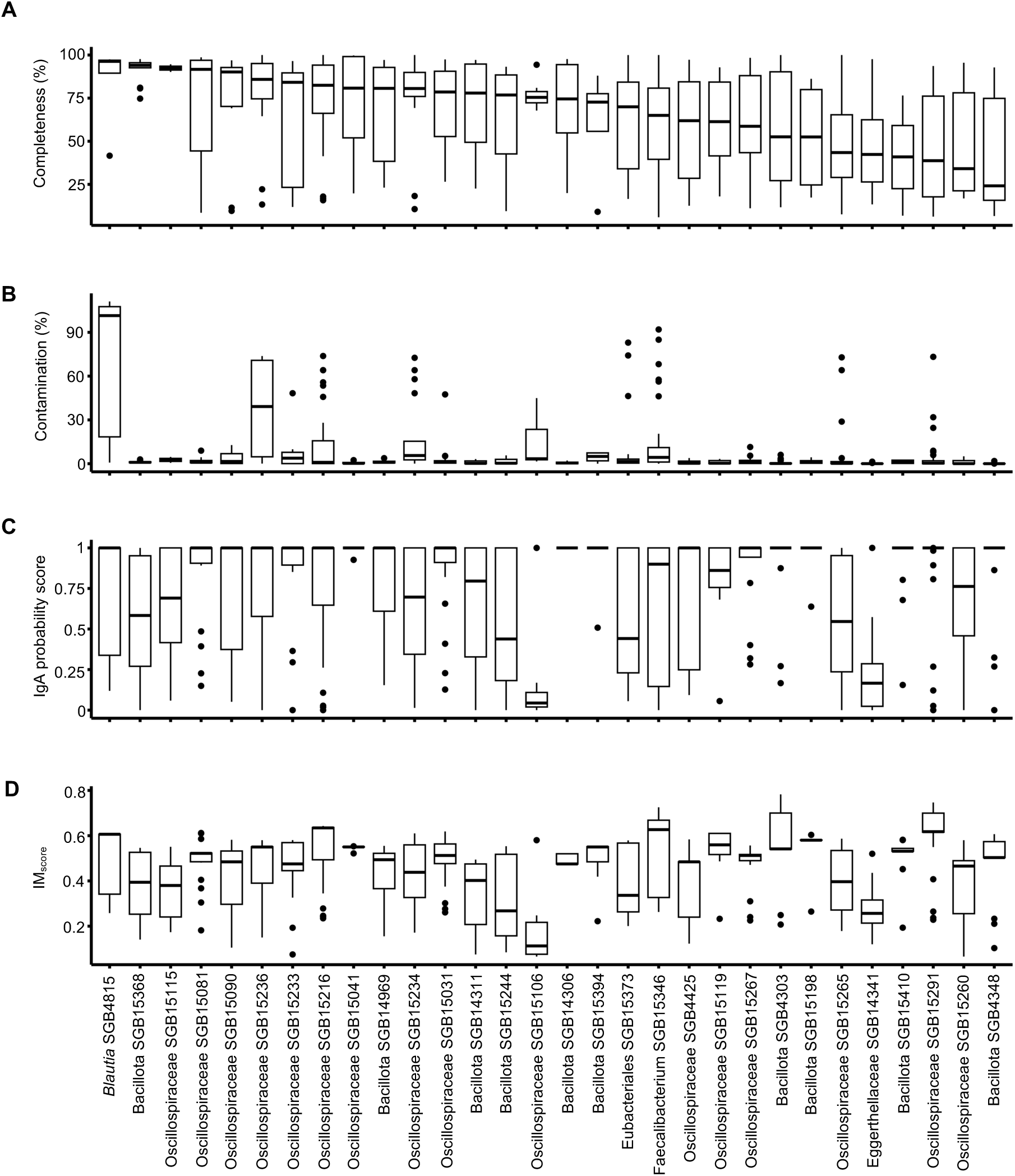
Yet-to-be-characterised IgA-bound microbial genomes reconstructed with high level of genome completeness, related to Figure 3. (A) Reconstructed MAGs’ completeness (A) based on metagenomics data of IgA+ fraction from this study, Olm et al. and van Gogh et al. study for yet-to-be-characterized MAGs with >50% prevalence in at least one study. (B) Low genome contamination is observed for for yet-to-be-characterised MAGs depicted in IgA-targeted microbiota from this study, Olm et al. and van Gogh et al. (C-D) IgA binding (C) and IM_score_ (D) for yet-to-be-characterised core microbiota members from Olm et al. and van Gogh et al.

**Figure S3.**
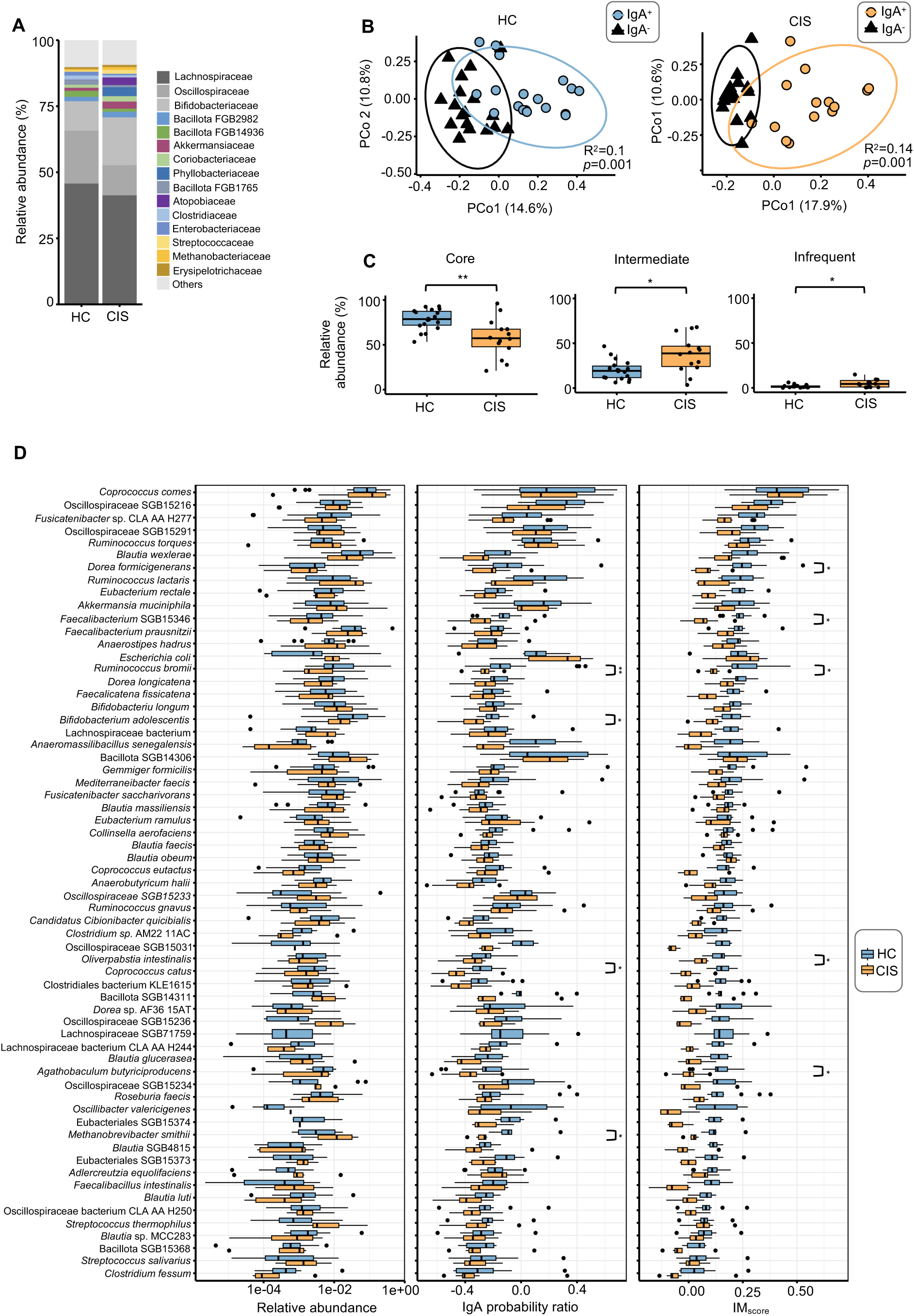
IgA-targeted gut microbiota in CIS subjects differs from healthy individuals, related to Figure 4. (A) Average relative abundance of the top 15 families of IgA-bound microbiota. (B). Beta-diversity using PCA analysis based on the Bray-Curtis dissimilarity for IgA+ and IgA-fractions for HC (left) and CIS (right). (C) Boxplots depicted species with different prevalence in HC and CIS patients. Species were classified based on the prevalence into core (prevalence >50%), intermediate (prevalence 10-50%) and infrequent taxa (prevalence <10%). (D) Differential abundance analysis of core microbiota for relative abundance of IgA-bound microbiota (left), IgA probability ratio (middle) and IM_score_ (right) between HC and CIS subjects. *p* values were from Wilcoxon rank-sum tests with BH correction. * p < 0.05, **p< 0.01, ***p<0.001, ****p<0.0001.

**Figure S4.**
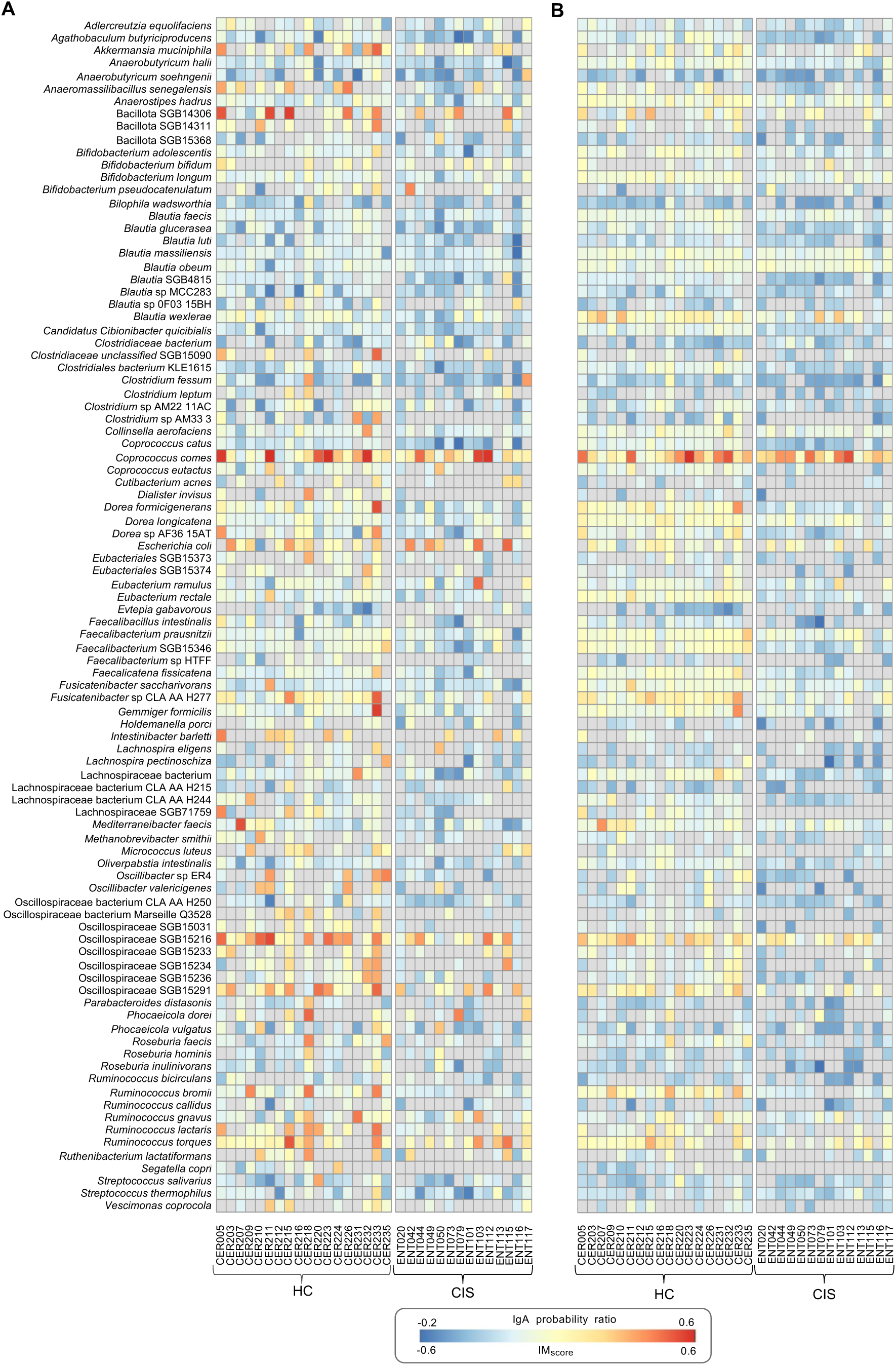
IgA-targeted microbiota at species level, related to Figure 4. (A-B) A heatmap representing IgA probability ratio (A) and IM_score_ (B) for top 100 prevalent species.

